# Dependence of the stimulus-driven microsaccade rate signature on visual stimulus polarity

**DOI:** 10.1101/2020.05.23.112417

**Authors:** Tatiana Malevich, Antimo Buonocore, Ziad M. Hafed

## Abstract

Microsaccades have a steady rate of occurrence during maintained gaze fixation, which gets transiently modulated by abrupt sensory stimuli. Such modulation, characterized by a rapid reduction in microsaccade frequency followed by a stronger rebound phase of high microsaccade rate, is often described as the microsaccadic rate signature, owing to its stereotyped nature. Here we investigated the impacts of stimulus polarity (luminance increments or luminance decrements relative to background luminance) and size on the microsaccadic rate signature. We presented brief visual flashes consisting of large or small white or black stimuli over an otherwise gray image background. Both large and small stimuli caused robust early microsaccadic inhibition, but only small ones caused a subsequent increase in microsaccade frequency above baseline microsaccade rate. Critically, small black stimuli were always associated with stronger modulations in microsaccade rate after stimulus onset than small white stimuli, particularly in the post-inhibition rebound phase of the microsaccadic rate signature. Because small stimuli were also associated with expected direction oscillations to and away from their locations of appearance, these stronger rate modulations in the rebound phase meant higher likelihoods of microsaccades opposite the black flash locations relative to the white flash locations. Our results demonstrate that the microsaccadic rate signature is sensitive to stimulus polarity, and they point to dissociable neural mechanisms underlying early microsaccadic inhibition after stimulus onset and later microsaccadic rate rebound at longer times thereafter. These results also demonstrate early access of oculomotor control circuitry to sensory representations, particularly for momentarily inhibiting saccade generation.

**New and noteworthy:** Microsaccades are small saccades that occur during gaze fixation. Microsaccade rate is transiently reduced after sudden stimulus onsets, and then strongly rebounds before returning to baseline. We explored the influence of stimulus polarity (black versus white) on this “rate signature”. We found that small black stimuli cause stronger microsaccadic modulations than white ones, but primarily in the rebound phase. This suggests dissociated neural mechanisms for microsaccadic inhibition and subsequent rebound in the microsaccadic rate signature.

## Introduction

Microsaccades occur occasionally during steady-state gaze fixation. When an unexpected stimulus onset occurs under such steady-state conditions, as is the case in a variety of behavioral experiments requiring maintained fixation (Hafed et al. 2015), stereotyped changes in microsaccade likelihood (and other properties) are known to take place. Specifically, microsaccade likelihood, or rate per second, abruptly decreases shortly after stimulus onset, remains near zero for a brief period of time, and then momentarily rebounds to higher rates than before stimulus onset (Bonneh et al. 2015; Buonocore et al. 2017a; Engbert and Kliegl 2003; Hafed and Ignashchenkova 2013; Laubrock et al. 2005; Peel et al. 2016; Rolfs et al. 2008; Tian et al. 2018; Valsecchi et al. 2007; White and Rolfs 2016). This pattern has been termed the “microsaccadic rate signature” (Engbert and Kliegl 2003; Hafed and Ignashchenkova 2013; Rolfs 2009; Rolfs et al. 2008; Scholes et al. 2015), owing to its highly repeatable nature across many paradigms, and it is also related to the more general phenomenon of saccadic inhibition reported for larger saccades (Bompas and Sumner 2011; Buonocore and McIntosh 2008; Buonocore et al. 2016; Edelman and Xu 2009; Reingold and Stampe 1999; 2004; 2002; 2003).

The neural mechanisms behind the microsaccadic rate signature, and saccadic inhibition in general, are still being investigated. Neurophysiological perturbation studies in the superior colliculus (SC), frontal eye fields (FEF), and primary visual cortex (V1) have resulted in initial informative steps towards clarifying these mechanisms. First, using a paradigm involving peripheral stimulus onsets, Hafed and colleagues demonstrated that monkeys exhibit the same microsaccadic rate signature as humans (Hafed et al. 2011). These effects persisted even after thousands of trials performed by the same animals in the same tasks, confirming the systematic nature of the effects. These authors then exploited the observation that monkeys exhibit the same phenomenon as humans to perform invasive neurophysiology; they reversibly inactivated portions of the SC topographic map representing the locations of the appearing peripheral stimuli (Hafed et al. 2013). The microsaccadic rate signature was virtually unaltered, whereas microsaccade directions were significantly redistributed (Hafed et al. 2013), consistent with a dissociation between the microsaccade rate signature and microsaccade direction oscillations after stimulus onsets (Buonocore et al. 2017a; Hafed and Ignashchenkova 2013; Tian et al. 2016). In follow up work, Peel and colleagues extended these results by reversibly inactivating the FEF. They found that the early inhibition was again unaltered, but, critically, the rebound phase of the microsaccadic rate signature was affected (Peel et al. 2016); there were fewer post-inhibition microsaccades than without FEF inactivation. In V1, lesions were found to affect microsaccades in general, but the early inhibition after stimulus onset was generally still present (Yoshida and Hafed 2017). Together with computational modeling (Hafed and Ignashchenkova 2013; Tian et al. 2016), all of these initial results suggest that there may be different components associated with the rate signature (e.g. inhibition versus rebound) that are mediated by distinct neural circuits; the early inhibition is clearly distinct from the later rebound that seems to particularly require frontal cortical control.

That said, the microsaccadic rate signature in its entirety must be related to early sensory responses, since the inhibition phase starts with very short latencies from stimulus onset (approximately 60-70 ms in monkeys) (Hafed et al. 2011; Malevich et al. 2020; Tian et al. 2018). It is, therefore, worthwhile to explore the effects of stimulus properties on subsequent microsaccadic modulations. For example, Rolfs and colleagues investigated the impacts of luminance and color contrast, as well as auditory stimulation, on microsaccadic inhibition (Rolfs et al. 2008; White and Rolfs 2016). Similarly, contrast sensitivity was related to the microsaccadic rate signature in other recent studies (Bonneh et al. 2015; Scholes et al. 2015). In all of these investigations, the general finding was that the strength of both inhibition and subsequent rebound increases with increasing stimulus strength. This suggests that expected sensory neuron properties (e.g. increased neural activity with increased stimulus contrast) must act rapidly on the oculomotor system to mediate inhibition, and potentially also influence subsequent rate rebounds. Here, we add to such existing descriptive studies about the microsaccadic rate signature. We document new evidence that visual stimulus polarity matters. We presented localized as well as diffuse visual flashes that were either white or black, relative to an otherwise gray background. We found that black localized stimuli were particularly effective in modulating the microsaccadic rate signature when compared to white stimuli, especially in the rebound phase, even when the white stimuli had higher contrast relative to the background.

Besides helping to clarify the properties of sensory pathways affecting the microsaccadic rate signature, our results are additionally important because of existing links between the rate signature and spatial attention shifts (Engbert and Kliegl 2003; Hafed 2013; Hafed et al. 2015; Hafed and Clark 2002; Tian et al. 2018; 2016). Despite accumulated evidence on differential effects of stimulus contrast on both so-called facilitatory and inhibitory cueing effects and on reaction times in general (Hawkins et al. 1988; Hughes 1984; Kean and Lambert 2003; Reuter-Lorenz et al. 1996), the question of whether and to what extent stimulus polarity itself affects cueing effects, to our knowledge, has not been explicitly addressed. This question might be of special interest, since “darks” / “blacks” seem to have temporal and sensitivity advantages over “whites” in visual perception, and there are perceptual asymmetries in processing of low and high luminances (Chubb and Nam 2000; Komban et al. 2011; Komban et al. 2014; Lu and Sperling 2012). Stimulus polarity can also activate distinct neural pathways as early as the retina through ON and OFF retinal image processing pathways (Chichilnisky and Kalmar 2002; Jin et al. 2011; Komban et al. 2011; Komban et al. 2014; Nichols et al. 2013; Xing et al. 2010; Yeh et al. 2009). Because we believe that microsaccades can potentially play an integral role in cognitive processes like covert attention (Chen et al. 2015; Hafed 2013; Hafed et al. 2015; Hafed and Clark 2002; Tian et al. 2018; 2016), we believe that knowing more about the stimulus conditions (and pathways) that might maximize or minimize the likelihood of microsaccades in a given paradigm would be useful in cognitive and systems neuroscience in general.

## Methods

### Ethics approvals

All monkey experiments were approved by ethics committees at the Regierungspräsidium Tübingen. The experiments were in line with the European Union directives and the German laws governing animal research.

### Laboratory setups

Monkey experiments were performed in the same laboratory environment as that described recently (Buonocore et al. 2019; Malevich et al. 2020; Skinner et al. 2019). A subset of the data (full-screen flash condition described below) were analyzed in brief in (Malevich et al. 2020), in order to compare the timing of microsaccadic inhibition to the novel ocular position drift phenomenon described in that study. However, the present study describes new analyses and comparisons to different stimulus conditions that are not reported on in the previous study.

The monkeys viewed stimuli on a cathode-ray-tube (CRT) display running at 120 Hz refresh rate. The display was gamma-corrected (linearized), and the stimuli were grayscale. Background and stimulus luminance values are described below with the behavioral tasks. Stimulus control was achieved using the Psychophysics Toolbox (Brainard 1997; Kleiner et al. 2017; Pelli 1997). The toolbox acted as a slave device receiving display update commands from a master device and sending back confirmation of display updates. The master system consisted of a real-time computer from National Instruments controlling all aspects of data acquisition (including digitization of eye position signals) and reward of the animals (in addition to display control). The real-time computer communicated with the Psychophysics Toolbox using direct Ethernet connections and universal data packet (UDP) protocols (Chen and Hafed 2013).

### Animal preparation

We collected behavioral data from 2 adult, male rhesus macaques (Macaca Mulatta). Monkeys M and A (aged 7 years, and weighing 9-10 kg) were implanted with a scleral coil in one eye to allow measuring eye movements (sampled at 1KHz) using the electromagnetic induction technique (Fuchs and Robinson 1966; Judge et al. 1980). The monkeys were also implanted with a head holder to stabilize their head during the experiments, with details on all implant surgeries provided earlier (Chen and Hafed 2013; Skinner et al. 2019). The monkeys were part of a larger neurophysiology project beyond the scope of the current manuscript.

### Monkey behavioral tasks

The monkeys maintained fixation on a small square spot of approximately 5 x 5 min arc dimensions. The spot was white (86 cd/m^2^) and drawn over a uniform gray background (29.7 cd/m^2^) in the rest of the display. The display subtended approximately +/- 15 deg horizontally and +/- 11 deg vertically relative to central fixation, and the rest of the laboratory setup beyond the display was dark. After approximately 550-1800 ms of initial fixation, a single-frame (~8 ms) flash occurred to modulate the microsaccadic rate signature. In different conditions, the flash could be either a full-screen flash, for which microsaccades were only partially analyzed in (Malevich et al. 2020), or a localized flash (not previously analyzed). The latter was a square of 1 x 1 deg dimensions centered on either 2.1 deg to the right or left of the fixation spot. On randomly interleaved control trials, the flash was sham (i.e. no flash was presented), and nothing happened on the display until trial end. Approximately 100-1400 ms after flash onset, the fixation spot disappeared, and the monkeys were rewarded for maintaining gaze fixation at the fixation spot throughout the trial. Note that this paradigm is the fixation variant of the paradigm that we used earlier during smooth pursuit eye movements generated by the same monkeys (Buonocore et al. 2019).

In one block of sessions, the stimuli used could be white flashes of the same luminance as the fixation spot (5167 trials analyzed from monkey M and 3104 trials analyzed from monkey A). In another block, the stimuli were all black flashes, but the fixation spot was still white (1513 trials analyzed from monkey M and 1818 trials analyzed from monkey A). Because we hypothesized that black flashes would have stronger influences in general than white flashes, motivated by earlier evidence in visual perception studies (Komban et al. 2011; Komban et al. 2014; Lu and Sperling 2012), we aimed to ensure that such stronger influences would be independent of stimulus contrast relative to the background. That is, because stimulus contrast can affect the microsaccadic rate signature (as detailed above in Introduction), we avoided a potential confound of stimulus contrast by having our background gray luminance level being closer to black than to white. Thus, relative to the background luminance, the contrast of black flashes was lower than that of white flashes. Yet, as we report in Results, black flashes often still had significantly stronger impacts on the microsaccadic rate signature, especially with the localized stimuli.

### Behavioral analyses

We detected microsaccades using established methods reported elsewhere (Bellet et al. 2019; Chen and Hafed 2013). Both methods rely on a mathematical differential (i.e. speed) or more (i.e. acceleration) of the digitized eye position signals acquired by our systems, with specific parameters for the classification of saccadic events depending on the specific signal noise levels in the digitized signals. We manually inspected each trial to correct for false alarms or misses by the automatic algorithms, which were rare. We also marked blinks or noise artifacts for later removal. In scleral eye coil data, blinks are easily discernible due to well-known blink-associated changes in eye position.

We estimated microsaccade rate as a function of time from stimulus onset using similar procedures to those we used earlier (Buonocore et al. 2017a; Hafed et al. 2011; Malevich et al. 2020). Briefly, for any time window of 80 ms duration and in any one trial, we counted how many microsaccades occurred within this window (typically 0 or 1). This gave us an estimate of instantaneous rate within such a window (i.e. expected number of microsaccades per window, divided by 80 ms window duration). We then moved the window in steps of 5 ms to obtain full time courses. The mean microsaccade rate curve across all trials of a given condition was then obtained by averaging the individual trial rate curves, and we obtained the standard error of the mean as an estimate of the dispersion of the across-trial measurements. Since some trials ended before 500 ms after flash onset (see *Monkey behavioral tasks* above), the across-trial average and standard error estimates that we obtained for any given time bin were restricted to only those individual trials that had data in this time bin; this was a majority of trials anyway. Also, because of the window duration and step size, the time courses were effectively low-pass filtered (smoothed) estimates of microsaccade rate (Bellet et al. 2017). We did not analyze potential higher frequency oscillations in microsaccade rate time courses. These tend to come later after the rebound phase anyway (Tian et al. 2016). We also confirmed that pre-stimulus baseline microsaccade rate in a given monkey was similar in the separate blocks of white and black flashes, therefore allowing us to compare and contrast polarity effects on the rate signature after flash onsets.

With localized flashes, we also considered microsaccade rate time courses independently for specific subsets of microsaccade directions. We specifically considered microsaccades that were either congruent or incongruent with flash location (meaning that we pooled right flash and left flash conditions together for these analyses). Congruent microsaccades were defined as those movements with a horizontal component in the direction of the flash. Incongruent microsaccades were defined as movements with a horizontal component opposite the flash location. Our past work shows that this categorization based on only the horizontal component of microsaccades is sufficient, since microsaccade vector directions after localized flashes are anyway highly systematically associated with the flash direction (Hafed and Ignashchenkova 2013; Tian et al. 2018). In related analyses, we also plotted direction distributions independently of microsaccade rate. Here, for every time bin relative to stimulus onset, we calculated the fraction of microsaccades occurring within this time bin that were congruent with flash location. This gave us a time course of direction distributions for all microsaccades that did occur (whether during the inhibition or rebound phases of the microsaccadic rate signature).

To analyze the time courses of microsaccade radial amplitudes after stimulus onset, we used similar procedures to the rate calculations described above. That is, we used a time window of 80 ms that was stepped in 5 ms steps to estimate the time courses of microsaccade amplitude modulations associated with different types of stimulus onsets in our experiments.

### Statistical analyses

All figures show and define error bars, which encompassed the standard error bounds around any given curve.

To statistically test the difference in the microsaccadic rate signature between conditions, we used non-parametric permutation tests with cluster-based correction for multiple comparisons (Maris and Oostenveld 2007), as we also described in detail in (Bellet et al. 2017; Idrees et al. 2020). First, for each time point (a bin) within an interval from −100 ms till +500 ms relative to stimulus onset, we compared two given conditions (e.g. localized versus full-screen flashes) by calculating the mean difference in their microsaccade rate. In order to obtain the null experimental distribution, we collected the trials from both conditions into a single set and, while maintaining the initial ratio of numbers of trials in each of the conditions, we randomly permuted their labels; we repeated this procedure 1000 times and recalculated the test statistic (i.e. the difference in rate curves between the two conditions) on each iteration. Second, we selected the bins of the original data whose test statistics were either below the 2.5^th^ percentile or above the 97.5^th^ percentile of the permutation distribution (i.e. significant within the 95% confidence level). For adjacent time bins having significant differences (i.e. for clusters of significance), we classified them into negative and positive clusters based on the sign of the difference in rate curves between the two conditions (i.e. clusters had either a negative or positive difference between the two compared microsaccade rate curves). We also repeated this procedure for each random permutation iteration by testing it against all other 999 random permutation iterations. This latter step gave us potential clusters of significance (positive or negative) that could arise by chance in the random permutations. Third, for both the observed and permuted data, we calculated the cluster-level summary statistic; this was defined as the sum of all absolute mean differences in any given potentially “significant” cluster. After that, we computed the Monte Carlo p-values of the original data’s clusters by assessing the probability of getting clusters with larger or equal cluster-level statistics under the null distribution (i.e. by taking the count of null data clusters with test statistics equal to or larger than the test statistic of any given original data cluster and dividing this count by the number of permutations that we used). A p-value of 0 indicated that none of the clusters of the null distribution had larger or equal cluster-level statistics than the real experimental data.

When testing either the localized or full-screen flash conditions against the control condition, the test was two-sided (i.e. looking for either positive or negative clusters) to avoid mutual masking of the expected inhibition and rebound effects. In this case, positive and negative clusters (i.e. clusters with positive and negative mean rate differences, respectively) in the experimental data were compared with positive and negative clusters in the permuted data, respectively; the clusters whose p-values exceeded the critical alpha level of 0.025 were considered as significant. All other comparisons were done with a one-tailed test, whereby the clusters were compared in their absolute value regardless of their sign; the critical alpha level was set to 0.05 in this case. The same algorithm was applied to the time course analyses of microsaccade amplitudes, except that here, all tests were one-sided.

When comparing magnitudes of the effects in different phases of the microsaccadic rate signature across conditions, we ran additional non-parametric permutation tests on the differences in minimum microsaccade rates during the inhibition phase or differences in peak microsaccade rates in the rebound phase, as well as in their latencies. To that end, based on the observations across monkeys, we predefined time intervals of interest for both microsaccadic inhibition (i.e. 70-180 ms after stimulus onset) and post-inhibition (i.e. 180-340 ms after stimulus onset) periods. For each experimental condition, we computed the mean microsaccade rate within such a predefined interval and found its extreme value (i.e. the minimum mean inhibition rate or the maximum mean rebound rate) and its latency relative to stimulus onset. Then, we calculated the difference in these values between two given conditions. In order to obtain the null experimental distribution, we did the same procedure as described above: we collected the trials from both conditions into a single data set and randomly permuted their labels, while keeping the initial ratio of the numbers of trials across conditions. We repeated this procedure 1000 times and, on each iteration, we recalculated the test statistics (i.e. the differences between the rate values and their latencies, when applicable). Finally, we computed the Monte Carlo p-values of the observed experimental differences by assessing the probability of getting the null-hypothesis test values at least as extreme as the observed experimental values. Significance was classified based on a critical alpha level of 0.05. This procedure also helped us to ensure that we did not miss any effect with the cluster-based permutation analyses due to different temporal dynamics of the inhibition and post-inhibition phases of the microsaccadic rate signature across conditions.

The same method was used for amplitude analyses, but this time we compared the maximum amplitude values in the predefined inhibition time window (i.e. 70-180 ms after stimulus onset) and the minimum amplitude values in the post-inhibition period (i.e. 180-340 ms after stimulus onset) when contrasting experimental and control conditions. In all other cases, the time window of interest was narrowed to +/-5 ms from the minimum microsaccade rate (for the inhibition period) or maximum microsaccade rate (for the rebound period) retrieved for a given condition, and the analysis was performed on the microsaccade amplitudes averaged over this time window.

To assess the effect of stimulus polarity on microsaccade directionality irrespective of the microsaccade rate, we compared the fractions of congruent microsaccades (i.e. the sum of microsaccades towards the flash divided by the sum of all microsaccades that occurred in a given time bin) over time between the black and white localized flashes. For this purpose, we used a bootstrapping procedure to obtain the estimates of their dispersion. In particular, we randomly resampled our data with replacement 1000 times and computed the fraction of congruent microsaccades for each sample. The central tendency measure and the estimate of its standard error were retrieved by calculating the mean and standard deviation of the bootstrap distribution.

Finally, when comparing fractions of congruent microsaccades or microsaccade amplitudes across conditions, we complemented the data visualization in the figures with microsaccade frequency histograms as a function of time, with bin widths of 24 ms and normalized with respect to the total number of trials in a given condition. This was done to provide an easier visual comparison between direction or amplitude effects and microsaccade rate. Such histograms are shown at the bottom of each panel in the corresponding figures; their scales are arbitrary with respect to the y-axis but kept proportional across conditions within a given monkey.

### Data availability

All data presented in this paper are stored in institute computers and are available upon reasonable request.

## Results

We documented the properties of the microsaccadic rate signature in two rhesus macaque monkeys as a function of either visual stimulus size - diffuse (full-screen flash condition) versus localized (localized flash condition) - or visual stimulus polarity - white versus black. In what follows, we first characterize the diffuse flash results before switching to the localized flash ones.

### Microsaccadic inhibition is similar for diffuse and localized visual flashes

Our full-screen flash condition created a diffuse stimulus over an extended range of the visual environment (approximately +/- 15 deg horizontally and +/- 11 deg vertically). On the other hand, our localized flash was much smaller (1 x 1 deg centered at 2.1 deg eccentricity). Both kinds of flashes were presented for only one display frame (~8 ms) over a uniform gray background filling the display (Methods); the rest of the laboratory was dark. We first asked whether microsaccadic inhibition would occur for both conditions, and whether it would exhibit different properties across them. For example, if microsaccadic inhibition is a function of sensory neuron properties (as alluded to in Introduction), then could surround suppression effects (Hubel and Wiesel 1968; Knierim and Van Essen 1992) associated with large, diffuse stimuli weaken or delay the occurrence of microsaccadic inhibition? If so, then this would implicate specific sensory areas, which are particularly sensitive to surround suppression effects, in contributing to the inhibition phase of the microsaccadic rate signature.

We plotted microsaccade rate as a function of time from stimulus onset for either diffuse or localized flashes (Methods). Figure 1A shows results with a localized black flash in monkey M, and Fig. 1B shows results with a diffuse (full-screen) black flash in the same monkey. In each panel, the gray curve shows microsaccade rate in the control condition in which no stimulus flash was presented (the two gray curves in the two panels are therefore identical). The red and blue horizontal bars on the x-axis of each plot show the significant clusters of time in which microsaccade rate was higher (red) or lower (blue) than control (cluster-based permutation tests; Methods). The results for the second monkey, A, are shown in Fig. 1D, E. In both monkeys, early microsaccadic inhibition occurred equally robustly regardless of whether the stimulus was diffuse or localized. That is, shortly after stimulus onset, there was a robust decrease in microsaccade likelihood before a subsequent rebound (compare colored to gray curves). The similarity of such decrease between the two stimulus types (localized versus diffuse) can be better appreciated by inspecting Fig. 1C, F, in which we plotted the microsaccade rate curves for the diffuse and localized flashes together on one graph. In monkey A, the initial microsaccadic inhibition phase was virtually identical with localized or diffuse black flashes (Fig. 1F); in monkey M, there was an earlier inhibition with the localized flash (Fig. 1C), but this effect was absent in the same monkey with white flashes instead of black ones, collected in separate blocks (Fig. 1G). Monkey A also had similar early inhibition profiles with white diffuse or white localized flashes (Fig. 1H).

**Figure 1.**
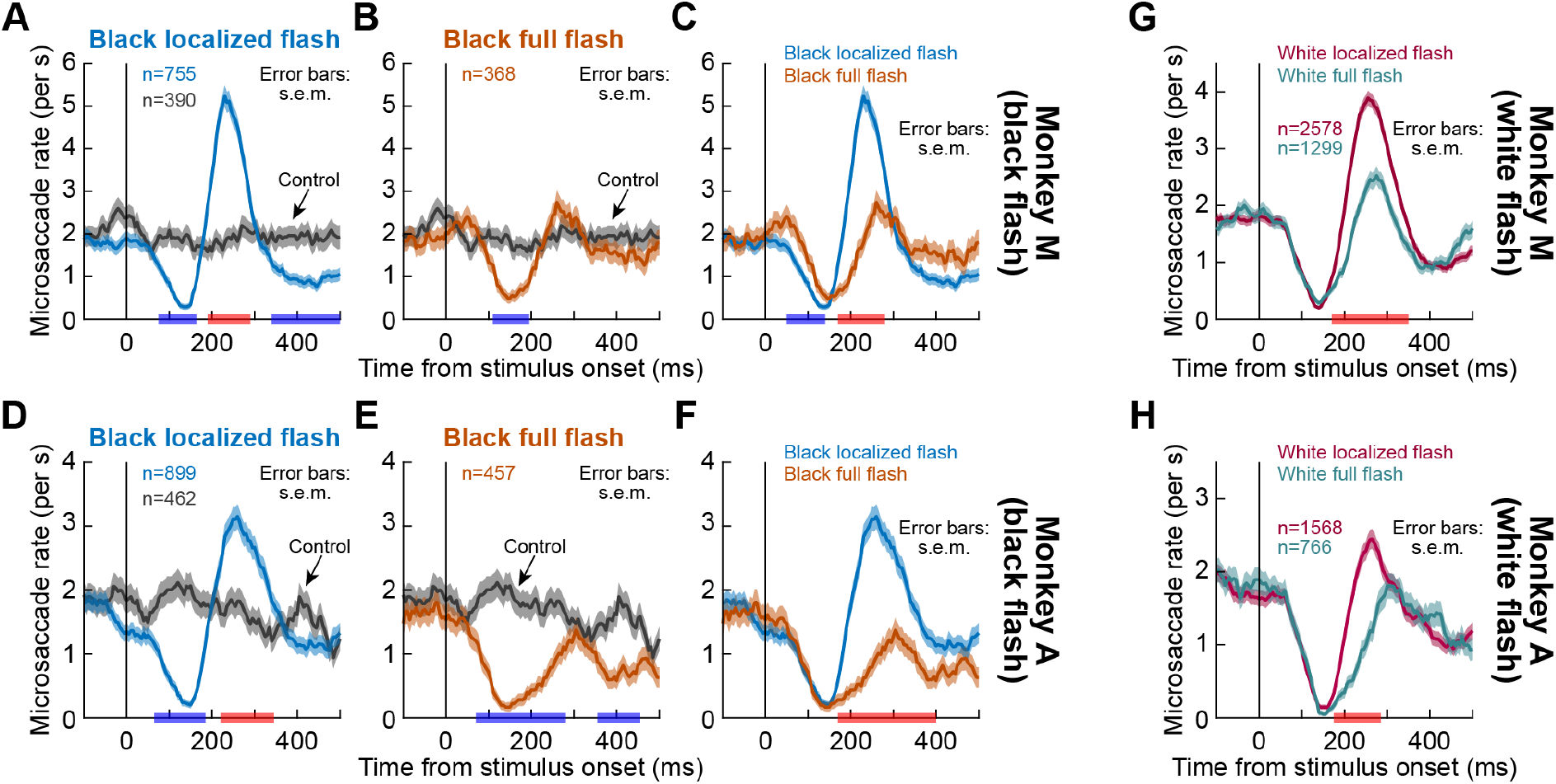
Microsaccade rate signatures with localized and diffuse visual stimuli. **(A)** Microsaccade rate in monkey M when a black localized flash appeared to the right or left of central fixation. The gray curve shows control microsaccade rate from trials in which the flash was absent. Relative to baseline control rates, microsaccade rate after flash onset decreased rapidly before rebounding. The rebound rate was higher than the control rate. At even longer intervals, microsaccade rate decreased again. Error bars denote s.e.m. bounds around each curve (Methods). The red and blue labels on the x-axis indicate positive (red) and negative (blue) significant clusters for the difference between conditions (flash minus control) (Methods). **(B)** Same data but when a full-screen flash was used. The early inhibition was similar to **A**, but the rebound was absent; microsaccade rate never went significantly above the control rate. **(C)** Microsaccadic rate signatures from **A**, **B** plotted together for easier comparison. Significance clusters on the x-axis now indicate whether the localized flash curve was higher (red) or lower (blue) than the full-screen flash curve. Significance in this case (i.e. the time points indicated on the x-axis) indicates that the two curves were different in absolute value regardless of the sign of the difference (Methods). **(D-F)** Same as **A**-**C**, but with monkey A data. Similar conclusions could be reached. **(G, H)** Same as **C**, **F**, but with white rather than black flashes (collected in separate blocks). Similar conclusions to **C**, **F** could be reached: inhibition occurred with both flash types, but rebound was stronger with localized flashes. Note that monkey M had earlier inhibition with localized than diffuse flashes only in the black flash condition; with white flashes, monkey M’s inhibition was similar for both flash types, like in monkey A.

Statistically, decreases in microsaccade rate started as early as 65-75 ms after stimulus onset in the localized black flash condition in both monkeys as well as in the diffuse black flash condition for monkey A. For monkey M, the inhibition was slightly delayed, starting at 110 ms after stimulus onset, with diffuse black flashes, as mentioned above. Specifically, for this monkey (M), the cluster-based permutation test that we used to investigate the properties of microsaccadic inhibition (Methods) revealed a rate difference between the localized and diffuse conditions during the interval 50-140 ms after stimulus onset (p = 0.017), consistent with a slightly later inhibition for diffuse flashes (Fig. 1C). Monkey A showed no difference in inhibition between localized and diffuse black flashes (Fig. 1F). In both monkeys, the time to peak inhibition was also not different across conditions (p = 0.243, 0.421 for monkeys M and A, respectively; black flashes). For white flashes, similar conclusions could be reached (p = 0.415, 0.277 for monkeys M and A, respectively).

In terms of the strength of microsaccadic inhibition, we measured microsaccade rate at the minimum after stimulus onset in the different conditions. We confirmed that localized and diffuse black flashes led to almost equally strong inhibitory effects as compared to the control condition in monkey M (mean minimum rate difference = - 1.318 microsaccades/s, p = 0 for localized flashes and mean minimum rate difference = −1.112 microsaccades/s, p = 0 for full-screen flashes). The difference between localized and full-screen flashes was not significant (p = 0.089). The measurements for monkey A were similar (mean minimum rate difference = −1.532 microsaccades/s, p = 0 for localized flashes and mean minimum rate difference = - 1.565 microsaccades/s, p = 0 for full-screen flashes), and the difference between localized and diffuse flashes was also not significant (p = 0.711). For white flashes, similar conclusions could be reached (p = 0.11, 0.073 for monkeys M and A, respectively, when comparing localized and diffuse flashes for minimum microsaccade rate).

Therefore, microsaccadic inhibition is equally strong with diffuse and localized visual stimuli. This adds to our earlier observations that even a simple luminance transient on the fixation spot itself is sufficient to induce strong microsaccadic inhibition (Buonocore et al. 2017a).

### Microsaccadic rate rebound is much weaker for diffuse than localized visual flashes, whether black or white

After the microsaccadic inhibition phase, there was a dramatic difference in the rebound phase of the microsaccadic rate signature between localized and diffuse flashes. In Fig. 1B, E, it can be seen that with full-screen flashes, post-inhibition microsaccade rate just returned to the baseline control rate without a clear “rebound” going above baseline. Targeted permutation tests revealed no difference in peak microsaccade rate (relative to control) in a predefined rebound interval (Methods) in monkey M (p = 0.098) and even showed an opposite effect in monkey A (mean peak rate difference = −0.489 microsaccades/s, p = 0.019). This is very different from how microsaccade rate strongly rebounded after the inhibition that was caused by localized flashes (Fig. 1A, D, indicated by red horizontal bars); peak rate was almost 3 times the baseline control rate in monkey M mean (peak rate difference = 3.04 microsaccades/s, p = 0; permutation test) and almost 2 times the baseline control rate in monkey A (mean peak rate difference = 1.288 microsaccades/s, p = 0; permutation test) (Fig. 1A, D; compare colored to gray curves).

We also compared the rate curves obtained with diffuse and localized flashes with each other by plotting them together (Fig. 1C, F). Cluster-based permutation tests revealed a significant difference between conditions in the rebound phase, starting at 170 ms after stimulus onset, for both monkeys (p = 0). As can be seen from Fig. 1C, F, peak microsaccade rate after the inhibition phase with localized flashes was more than 2 times stronger than peak microsaccade rate after the inhibition phase with diffuse flashes in both monkeys. We quantified these effects by running permutation tests on the peak rate values and their latencies. In monkey M, the mean peak rate difference between localized and diffuse flashes was 2.491 microsaccades/s (p = 0), and the latency difference was −30 ms (p = 0). These values were 1.777 microsaccades/s (p = 0) and −45 ms (p = 0.026), respectively, for monkey A.

With white flashes, similar conclusions could also be reached (Fig. 1G, H). In this case, significant differences between diffuse and localized conditions in the post-inhibition period emerged 170-175 ms after stimulus onset. Moreover, once again, with localized flashes, microsaccade rate reached its peak earlier (latency difference = −20 ms, p = 0.003 for monkey M and latency difference = −45 ms, p = 0.014 for monkey A; permutation tests) and rose higher (mean peak rate difference = 1.36 microsaccades/s, p = 0 for monkey M and mean peak rate difference = 0.643 microsaccades/s, p = 0.004 for monkey A; permutation tests) than with diffuse stimuli. However, note that the peak in microsaccade rate after localized flashes was notably lower than that with black flashes, as we describe in more detail later.

To further clarify whether the lack of post-inhibition microsaccadic rebound with diffuse flashes depended on stimulus polarity, we also plotted the white and black diffuse flash curves together and statistically assessed the difference between them (Fig. 2). There were again no apparent differences in time courses associated with stimulus polarity for diffuse flashes, except for a very late effect in the post-rebound interval in monkey A indicated by the blue bar in Fig. 2B. For both monkeys, stimulus polarity did not affect the peak rebound rate (p = 0.502 for monkey M and p = 0.093 for monkey A; permutation tests) nor its latency (p = 0.19 for monkey M and p = 0.429 for monkey A; permutation tests). Monkey A did show an earlier maximum inhibition for black diffuse flashes than for the white ones (latency difference = −10 ms, p = 0.015; permutation test) but no difference in its strength (p = 0.102; permutation test); neither of the effects reached significance in monkey M (p = 0.062 for minimum rate and p = 0.271 for latency; permutation tests).

**Figure 2.**
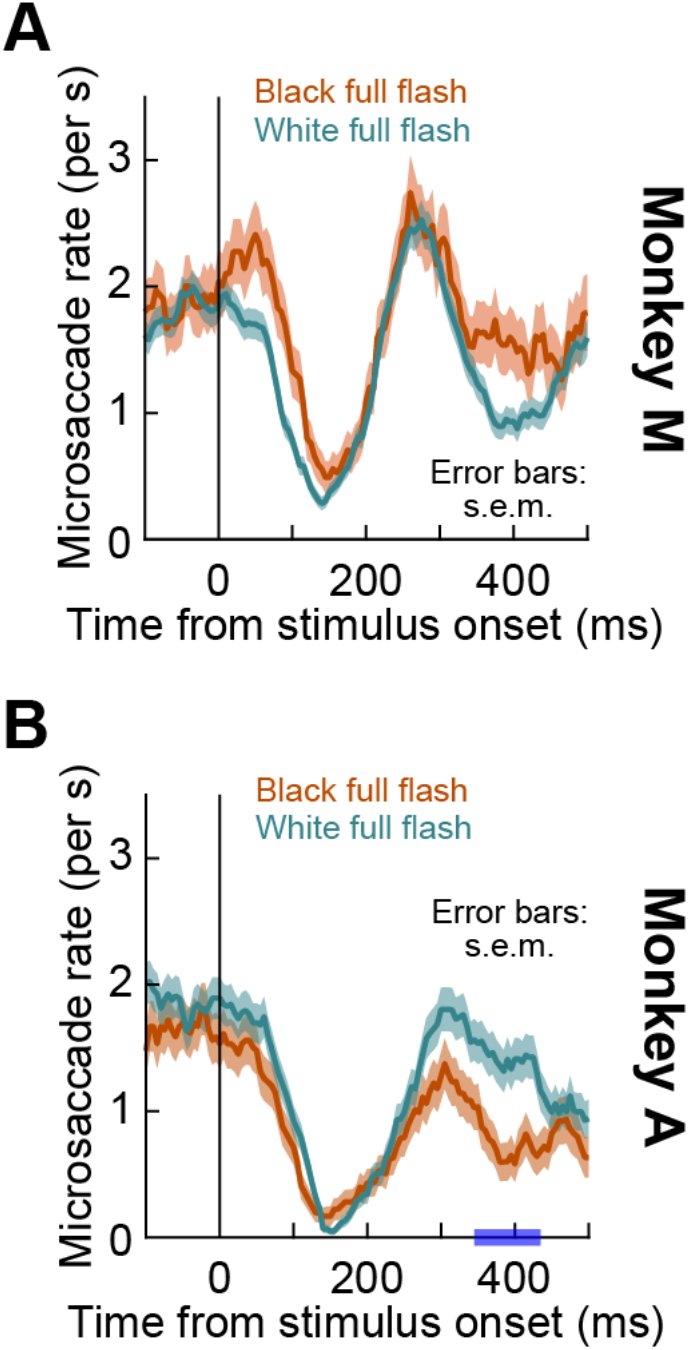
Microsaccade rate signatures with black and white diffuse visual stimuli. For each monkey, we plotted microsaccade rates from Fig. 1, this time directly comparing black versus white full-screen flashes. In monkey M, there was only a trend for stronger and earlier inhibition immediately after stimulus onset with white, rather than black, full-screen flashes (**A**). In monkey A, maximal inhibition was reached 10 ms earlier for black than white flashes, and there was only a trend for a stronger post-inhibition rebound in microsaccade rate with white, rather than black, full-screen flashes (**B**). Monkey A showed a later difference (blue interval on the x-axis delineating a significant interval of negative mean difference between microsaccade rates in the black and white diffuse flash conditions, obtained with the one-sided cluster-based permutation test; p = 0.002; Methods). All other conventions are similar to Fig. 1.

The above results, so far, suggest that diffuse visual stimuli are as effective as localized visual stimuli in causing robust microsaccadic inhibition in rhesus macaque monkeys (Fig. 1). However, post-inhibition microsaccade rates are much lower with diffuse stimuli (Fig. 1). Moreover, these effects with diffuse stimuli are largely independent of stimulus polarity (Fig. 2). There were also no clear effects on microsaccade direction distributions with diffuse stimuli (data not shown), as might be expected due to the symmetric nature of the full-screen flashes relative to the fixation spot. We next explored the localized stimulus conditions in more detail, highlighting a significant difference in microsaccadic rate signatures as a function of stimulus polarity.

### Black localized flashes have stronger “cueing effects” than white localized flashes

With localized flashes, we saw above that the microsaccadic rate signature looked more similar to classic literature descriptions. That is, there was a strong post-inhibition rebound in microsaccade rate, reaching levels significantly higher than baseline microsaccade-rate during steady-state fixation (colored versus gray curves in Fig. 1A, D, indicated by red horizontal bars on the x-axes). However, comparing the different y-axis scales used in Fig. 1C, F and Fig. 1G, H additionally revealed an influence of stimulus polarity. Unlike in Fig. 2, there was a substantial effect of black flashes in particular on the microsaccadic rate signature with localized stimulus onsets. This effect can be seen clearly in Fig. 3; black localized flashes were particularly effective in modulating the post-inhibition rebound phase of the microsaccadic rate signature, as was also confirmed by cluster-based permutation tests (the red horizontal bars on the x-axes in Fig. 3 indicate the regions of significantly stronger rebound with black flashes; p = 0 for both monkeys). Both monkeys showed a significantly higher peak in microsaccade rate with black, rather than white, visual stimuli (mean peak rate difference = 1.344 microsaccades/s, p = 0 for monkey M and mean peak rate difference = 0.705 microsaccades/s, p = 0.002 for monkey A; permutation tests). In addition, the rate reached its maximum 25 ms faster with the black stimuli in the case of monkey M, whereas monkey A showed a similar, albeit not significant, trend (p = 0.001 for monkey M and p = 0.152 for monkey A; permutation tests). These observations cannot be explained by stimulus contrast, because the contrast of the black flash relative to the background luminance was actually lower, by experimental design, than the contrast of the white flash relative to the background luminance (Methods).

**Figure 3.**
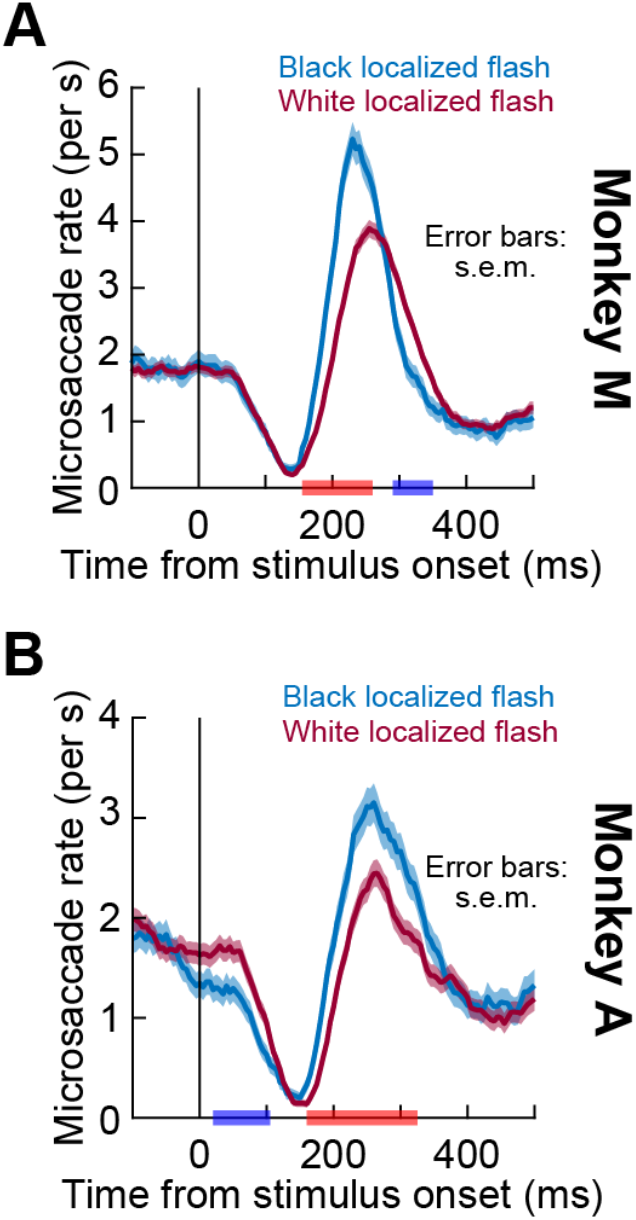
Microsaccade rate signatures with black and white localized visual stimuli. For each monkey, we plotted microsaccade rates from Fig. 1, this time directly comparing black versus white localized flashes, and we performed their time-course analyses with the cluster-based permutation tests described in Methods. The red and blue labels on the x-axes indicate significant intervals of positive and negative mean differences, respectively, between microsaccade rates in the black and white localized flash conditions, obtained with a critical alpha level of 0.05 (i.e. one-sided tests; Methods). In both monkeys, the post-inhibition microsaccadic rebound was significantly stronger with black than white localized flashes. This is different from the effects of stimulus polarity that we saw with diffuse flashes (Fig. 2). All other conventions are similar to Fig. 1.

In terms of the initial microsaccadic inhibition phase, it was generally similar whether black or white localized flashes were used. In monkey M, neither the cluster-based permutation test nor the analysis of the minimum inhibition rate or its latency brought significant results (p = 0.194 for minimum rate and p = 0.261 for latency; permutation tests). In monkey A, there was a significantly earlier inhibition effect caused by black localized flashes in the interval of 20-105 ms after stimulus onset (p = 0.023, cluster-based permutation test), which reached its maximum 10 ms faster than in the case of white flashes (p = 0.034; permutation test). However, the difference in the minimum inhibition rate was again not significant (p = 0.315; permutation test), which is in line with this monkey’s polarity effect in the inhibition period for the diffuse flashes.

Because localized visual stimuli have a directional component associated with them, they resemble “cues” in classic attentional cueing tasks. Past work has shown how such cues, even when task irrelevant (Buonocore et al. 2017a; Hafed and Ignashchenkova 2013), are associated with very systematic directional modulations of microsaccades when they appear under steady-state fixation conditions. When viewed from the perspective of the microsaccadic rate signature, these direction modulations consist of two primary effects: (1) a later inhibition of microsaccades that are congruent (in their direction) with stimulus location when compared to the inhibition time of microsaccades that are incongruent with stimulus location; and (2) a stronger and earlier post-inhibition rebound for microsaccades that are incongruent with stimulus location than for congruent microsaccades (Hafed et al. 2015; Hafed and Ignashchenkova 2013; Laubrock et al. 2005; Tian et al. 2018; 2016). In other words, microsaccades that do occur early after stimulus onset tend to be strongly biased towards the stimulus location, and microsaccades occurring late after stimulus onset tend to be biased in the opposite direction, and this is believed to reflect an interaction between ongoing microsaccade motor commands and visual bursts associated with stimulus onsets (Buonocore et al. 2017a; Hafed and Ignashchenkova 2013). When we analyzed microsaccadic rate signatures for different microsaccade directions in our localized flash conditions, we confirmed these expected results, although the later rebound effect was weaker in monkey A in general. Critically, the effects were always stronger with black than white stimuli in both monkeys, consistent with Fig. 3.

Specifically, in Fig. 4A, we plotted the rate of congruent and incongruent microsaccades with a localized black flash for monkey M. Congruent microsaccades were defined as those with directions towards the stimulus location, and incongruent ones were opposite the stimulus location (Methods). Congruent microsaccades were harder to inhibit than incongruent microsaccades (left black arrow), suggesting that in these early times after stimulus onset, if a microsaccade were to occur, it was more likely to be directed towards the flash location (Buonocore et al. 2017a; Hafed and Ignashchenkova 2013; Tian et al. 2018; 2016). Quantitatively, the rate curves near inhibition onset were different in the interval 35-140 ms after stimulus onset (p = 0.01; cluster-based permutation test). Moreover, maximal inhibition was statistically stronger for incongruent microsaccades (p = 0; permutation test), and the maximal inhibition latency was 20 ms earlier (p = 0.04). In the post-inhibition phase, the rate of incongruent microsaccades rose earlier and reached higher peaks than the rate of congruent microsaccades (peak rate difference, in absolute value, between the two curves = 2.484 microsaccades/s and peak latency difference = 10 ms; p = 0 and 0.005, respectively; permutation tests). The two curves started deviating from each other at 170 ms after stimulus onset (p = 0; cluster-based permutation test). These observations are consistent with the idea that later microsaccades were biased away from the stimulus location (Buonocore et al. 2017a; Hafed and Ignashchenkova 2013; Tian et al. 2018; 2016). Importantly, both effects were significantly weaker with white flashes (Fig. 4B). That is, the difference between the congruent and incongruent curves was smaller overall than in Fig. 4A, and the post-inhibition rebound rate was also weaker. In fact, with white flashes, the early difference in inhibition between congruent and incongruent microsaccades was virtually absent (Fig. 4B). Similarly, the difference in maximal microsaccade rebound rate was now 1.242 microsaccades/s (p = 0) as opposed to 2.484 microsaccades/s with black flashes.

**Figure 4.**
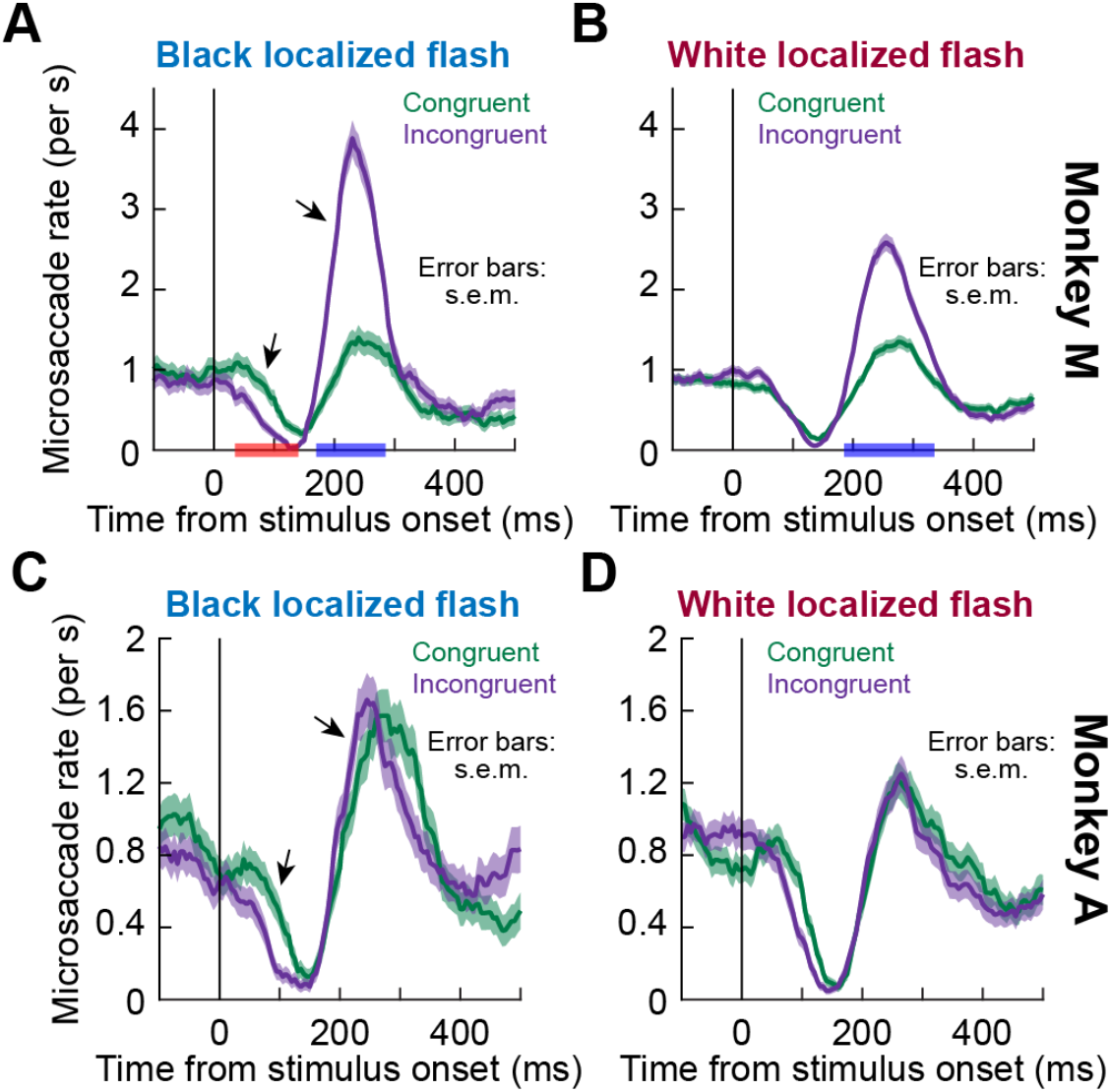
Microsaccade rate signatures with black and white localized visual stimuli when separated based on microsaccade direction. **(A)** Microsaccade rate in monkey M computed separately for congruent and incongruent microsaccades. Congruent microsaccades were defined as those movements directed towards the flash location, and incongruent microsaccades were defined as the microsaccades directed opposite the flash location (Methods). This panel shows results with a black flash. Consistent with earlier results, congruent microsaccades were harder to inhibit than incongruent microsaccades (left black arrow). Later in time, incongruent microsaccades were easier to generate than congruent microsaccades in the post-inhibition phase, as evidenced by the earlier and higher post-inhibition rise in microsaccade rate (right black arrow). **(B)** With white localized flashes, the difference between the congruent and incongruent curves was smaller overall than in **A**, both in the early inhibition phase as well as in the later rebound phase. Moreover, the overall rebound peak rate was lower than the peak rate with black flashes in **A**. See text for statistics. **(C, D)** Same results for monkey A. This monkey showed weaker effects than monkey M, but they were all consistent with the monkey M observations. That is, early directional differences associated with microsaccadic inhibition were weaker with white flashes (**D**) but amplified with black flashes (**C**; left black arrow). Moreover, post-inhibition rebound was slightly stronger for incongruent microsaccades with black (**C**; right black arrow) than white (**D**) localized flashes. All other conventions are similar to Fig. 1. Red and blue bars on x-axes show significant clusters of positive (red) and negative (blue) mean differences at the critical alpha level of 0.05 (Methods).

With monkey A, all of the effects described above were significantly weaker overall (Fig. 4C, D). Nonetheless, consistent with monkey M, black localized flashes were always associated with stronger trends (Fig. 4C; also see Fig. 5 below for further statistical comparisons). Closer inspection of this monkey’s eye movement data revealed a very strong bias to generate leftward microsaccades, even in baseline without any flashes. This strong bias masked the cueing effects after flash onset, which were still present but muted due to the large baseline directional bias. It is intriguing that even for a monkey like this one, for whom the “cueing effects” with white flashes were weak (Fig. 4D), they were still amplified with black flashes (Fig. 4C).

**Figure 5.**
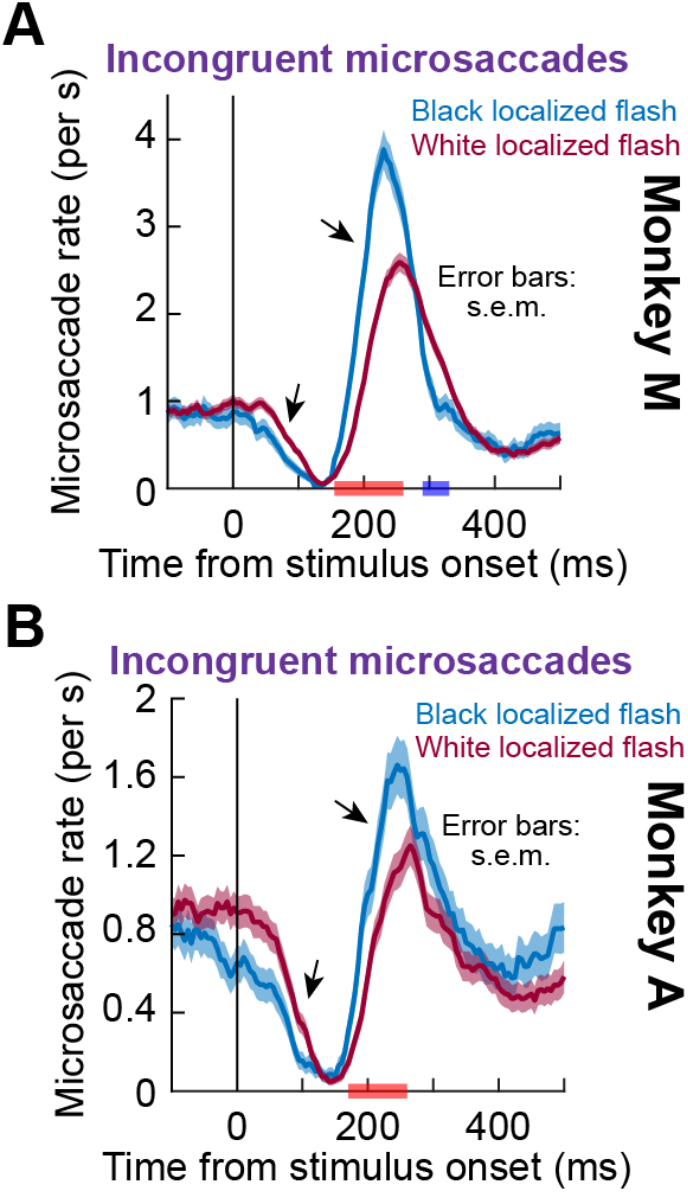
Effects of black localized flashes on incongruent microsaccades. Same data as in Fig. 4, but now showing the incongruent curves (purple in Fig. 4) for a given monkey under either white (dark pink) or black (blue) localized flash conditions. In both monkeys, black flashes were associated with higher rebound of incongruent microsaccades after the initial microsaccadic inhibition than white flashes (right black arrow in each panel). In both monkeys, there was also a trend for earlier microsaccadic inhibition time with black than with white flashes (left black arrow in each panel). The red horizontal bars on the x-axes denote significant clusters of positive mean differences between the black and white localized flash conditions (i.e. an earlier and stronger effect under the black flash condition, obtained with one-tailed cluster-based permutation tests at the critical alpha level of 0.05). The blue horizontal bar in **A** indicates a significant negative cluster showing an inverted pattern at the end of the rebound phase in monkey M. All other conventions are similar to Fig. 1.

Therefore, not only were black localized flashes associated with stronger microsaccadic rate modulations in both monkeys (Fig. 3), these stronger effects had a directional component, the largest of which was on enhancing the post-inhibition rebound of incongruent microsaccades (Fig. 4). So-called cueing effects on microsaccades were, thus, stronger with black than white localized flashes (at least in monkey M), even though the contrast of the black flashes relative to background luminance was lower.

To further explore this incongruent microsaccade effect in more detail, and to confirm that it was still present in monkey A despite the baseline directional bias alluded to above, we plotted the rates of only incongruent microsaccades, now separated based on whether the localized flash was white or black (Fig. 5). In both monkeys, the post-inhibition rate of incongruent microsaccades was significantly higher (and rose earlier) with black localized flashes than with white localized flashes (Fig. 5; right black arrow in each panel), as also revealed by cluster-based permutation tests (p = 0 for monkey M and p = 0.003 for monkey A; intervals: 155-260 ms and 170-260 ms in monkeys M and A, respectively). Also, both monkeys showed a stronger and earlier peak rate in the black flash condition than in the white flash condition (monkey M: peak rate difference = 1.293 microsaccades/s, p = 0 and latency difference = −25 ms, p = 0.001; monkey A: peak rate difference = 0.411 microsaccades/s, p = 0.015 and latency difference = −20 ms, p = 0.028; permutation tests). Interestingly, in both monkeys, there was a trend for earlier inhibition of incongruent microsaccades with black flashes when compared to white flashes (Fig. 5; left black arrow in each panel), which can explain the stronger cueing effects in Fig. 4A, C with black flashes. In monkey M, the inhibition of incongruent microsaccades with black flashes reached its maximum 10 ms earlier than with white flashes, although the difference in strength of the maximum inhibition was not significant (p = 0.023 for latency and p = 0.253 for minimum rate; permutation test). There were also no significant differences for the minimum rate and its latency in monkey A (p = 0.575 for minimum rate and p = 0.57 for latency; permutation tests).

For completeness, we next assessed microsaccade directions independently of microsaccade rates. For each time bin relative to localized flash onset time, we computed the fraction of microsaccades that both occurred within this time bin and were also congruent with flash location. This gave us a time course of microsaccade directions relative to the flash location, which we statistically assessed by performing bootstrapping with resampling (Methods). We did this separately for black and white flashes. These results are shown in Fig. 6, in which we also superimposed histograms of all microsaccade times in each flash condition in order to visually relate the microsaccade direction time courses with the microsaccadic rate signatures (the histograms in Fig. 6 are essentially another way to visualize the same rate curves of localized flashes in Fig. 1). As can be seen, in both monkeys, the likelihood of getting a microsaccade directed towards the flash sharply increased after stimulus onset, peaking at the time of maximal inhibition, which is consistent with previous findings (Buonocore et al. 2017a; Hafed and Ignashchenkova 2013; Pastukhov and Braun 2010; Tian et al. 2018; 2016). In the post-inhibition period, this pattern started to reverse, although only monkey M demonstrated a strong and expected bias in the direction opposite to the flash location (Buonocore et al. 2017a; Hafed and Ignashchenkova 2013; Tian et al. 2018; 2016). This is due to a strong bias in monkey A to make leftward microsaccades even in baseline, as mentioned above, which masked the transient flash effects. Nonetheless, both monkeys showed a trend for increased early bias in microsaccade directions towards flash location with black stimuli, consistent with Figs. 4, 5 above. In monkey M, the black flashes were also associated with more opposite microsaccades in the rate rebound phase after microsaccadic inhibition when compared to the white flashes. These results are again consistent with Figs. 4, 5 above.

**Figure 6.**
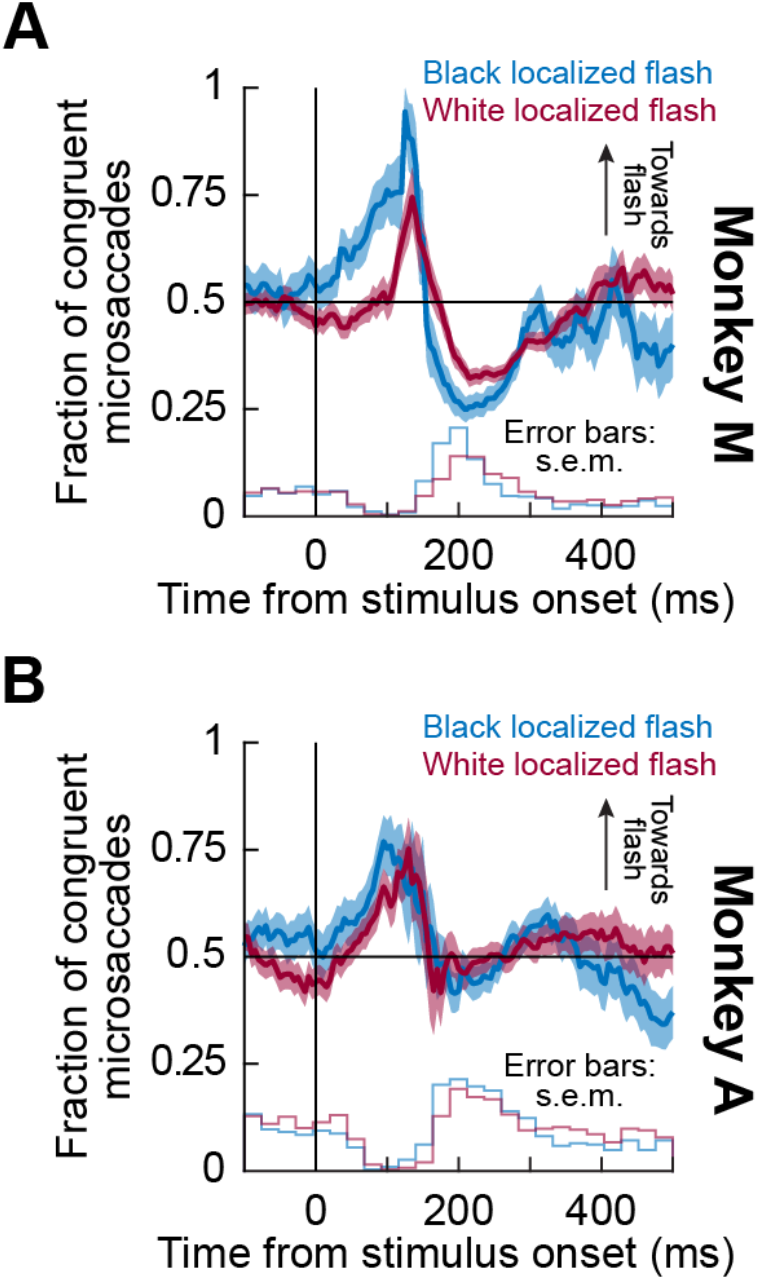
Distribution of microsaccade directions relative to localized flash location for black and white flashes. The thick curves with error bars show the time courses of fractions of microsaccades directed towards the flash location under either white or black localized flash conditions. The means and their standard errors were computed using bootstrapping with replacement. The histograms at the bottom of each panel show in the corresponding color the frequency of all microsaccades, regardless of their direction, that happened under the black and white localized flash conditions. The histograms were normalized with respect to the number of trials in a given condition; their scales are arbitrary with respect to the y-axis but kept proportional to each other within a given monkey. In both monkeys, the fraction of congruent microsaccades increased during the inhibition phase and started to decrease at the beginning of the rebound period. In addition, both monkeys showed a tendency for an earlier inhibition of incongruent microsaccades with black stimuli (i.e. stronger directional modulation towards the flash location), whereas only monkey M demonstrated an effect of stimulus polarity in the rebound phase after microsaccadic inhibition. All other conventions are similar to Fig. 1.

### Microsaccade amplitudes exhibit stronger temporal oscillations after stimulus onset with localized than diffuse flashes, but with no dependence on stimulus polarity

Because cue onsets, particularly when localized, can transiently modulate instantaneous foveal eye position errors at the fixation spot (Tian et al. 2018; 2016), microsaccade amplitude is also expected to be affected along with the microsaccadic rate signature (Buonocore et al. 2017a). We therefore documented the time courses of microsaccade amplitude variations after stimulus onset for our different stimulus sizes and polarities.

In terms of stimulus size, Fig. 7A-F shows comparisons between microsaccade amplitude time courses for diffuse and localized black flashes, similar in approach to Fig. 1A-F. In both monkeys, localized flashes caused marked modulations of microsaccade amplitude relative to baseline control amplitudes (Fig. 7A, D). Specifically, in the early inhibition phase in which microsaccades were likely to be directed towards the flash location (Fig. 4), microsaccade amplitude increased: permutation tests in the predefined period of 70-180 ms after stimulus onset (Methods) showed that peak microsaccade amplitudes for localized flashes were 0.156 deg (p = 0.032) higher in monkey M and 0.105 deg (p = 0.004) higher in monkey A than in control. So, the few microsaccades that did occur during microsaccadic inhibition were enlarged (Buonocore et al. 2017a). For later post-inhibition microsaccades, which were predominantly incongruent microsaccades (Figs. 4–6), microsaccade amplitude decreased: minimum amplitudes in the predefined interval of 180-340 ms after stimulus onset were 0.138 deg smaller than for the control in monkey M and 0.058 deg smaller than for the control in monkey A and (p = 0 and 0.016 for monkeys M and A, respectively; permutations tests). With full-screen flashes, both monkeys had a significant increase in the peak amplitudes in the early inhibition interval of 70-180 ms (mean difference = 0.257 deg, p = 0 for monkey M and mean difference = 0.109 deg, p = 0.003 for monkey A; permutation tests). However, there were no clear differences between microsaccade amplitudes with or without diffuse flashes later on in the post-inhibition period (compare colored to gray curves), even between minimum amplitudes within the predefined rebound interval (p = 0.723 for monkey M and p = 0.318 for monkey A; permutation tests). Moreover, direct comparisons between localized and diffuse flashes confirmed that rebound microsaccades became smaller in amplitude with localized, but not diffuse, flashes (Fig. 7C, F). Similar observations were made with white flashes (Fig. 7G, H). In fact, direct evaluation of stimulus polarities under the different stimulus size conditions showed that amplitude effects did not strongly depend on stimulus polarity (Fig. 8). If anything, there was a trend for white flashes, small or large, to be associated with stronger overall amplitude modulations as a function of time after stimulus onset (Fig. 8).

**Figure 7.**
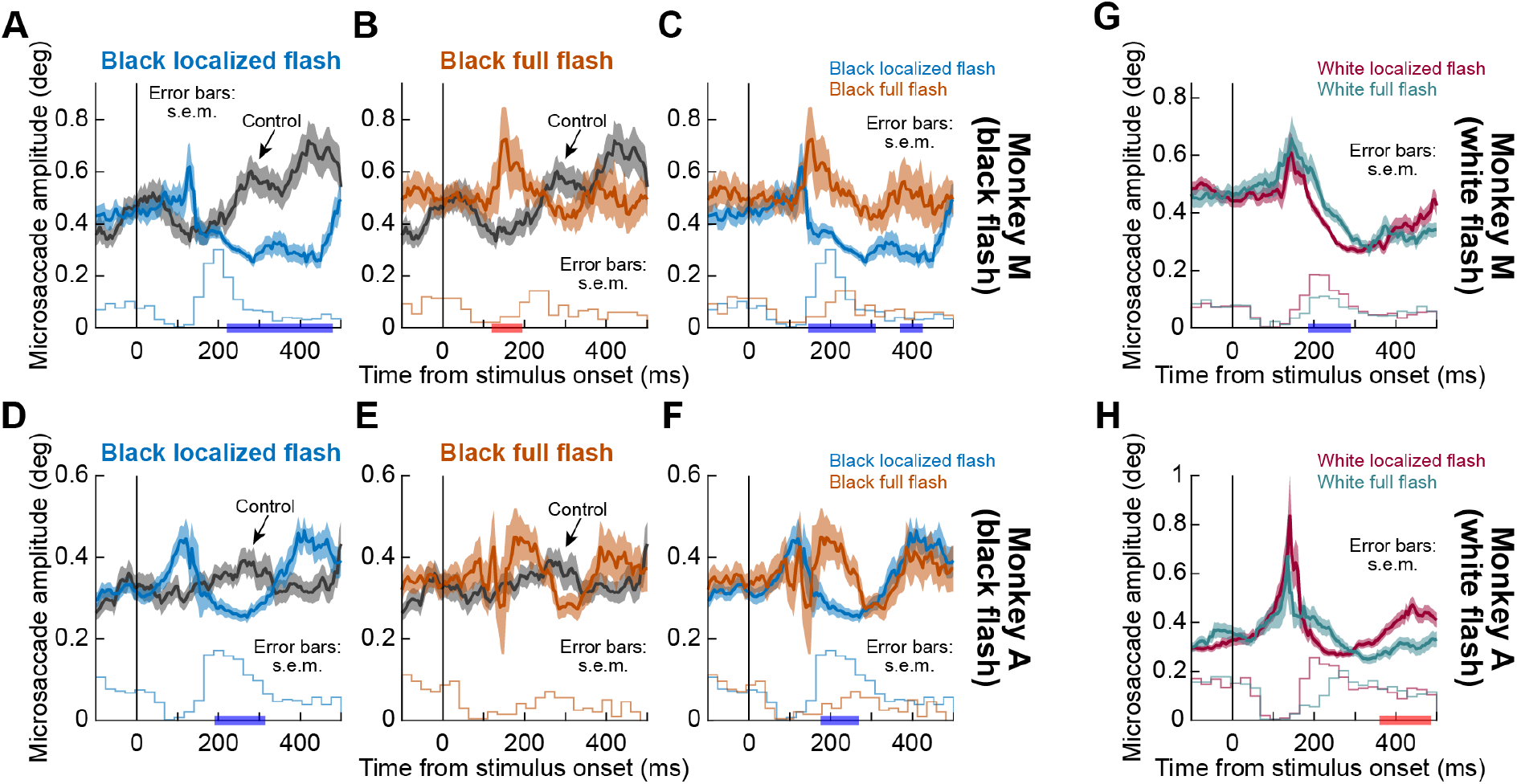
Microsaccade amplitudes for the data of Fig. 1. Same analyses as in Fig. 1, but for microsaccade amplitude. Localized black flashes (**A**, **D**) were associated with an initial amplitude increase during the initial inhibition phase of the microsaccadic rate signature followed by an amplitude decrease (relative to control) during the rebound phase. Full-screen flashes (**B**, **E**) did not show a clear decrease in amplitude in the post-inhibition phase of the microsaccadic rate signature. Direct comparisons between localized and diffuse black flashes are shown in (**C**, **F**), confirming that localized flashes caused stronger amplitude modulations. The effects with white flashes are shown in (**G**, **H**). Red and blue labels on x-axes indicate, respectively, clusters with positive and negative mean differences between localized (**A,D**) or full-screen (**B,E**) flashes and the control condition; or between localized and full-screen flashes (**C, F-H**). Significance was defined at the 0.05 level after cluster-based correction for multiple comparisons (Methods). The histograms at the bottom of each panel show, in the corresponding color, the frequency of microsaccades that happened under a given condition. The histogram conventions are the same as in Fig. 6. All other conventions are similar to Fig. 1.

**Figure 8.**
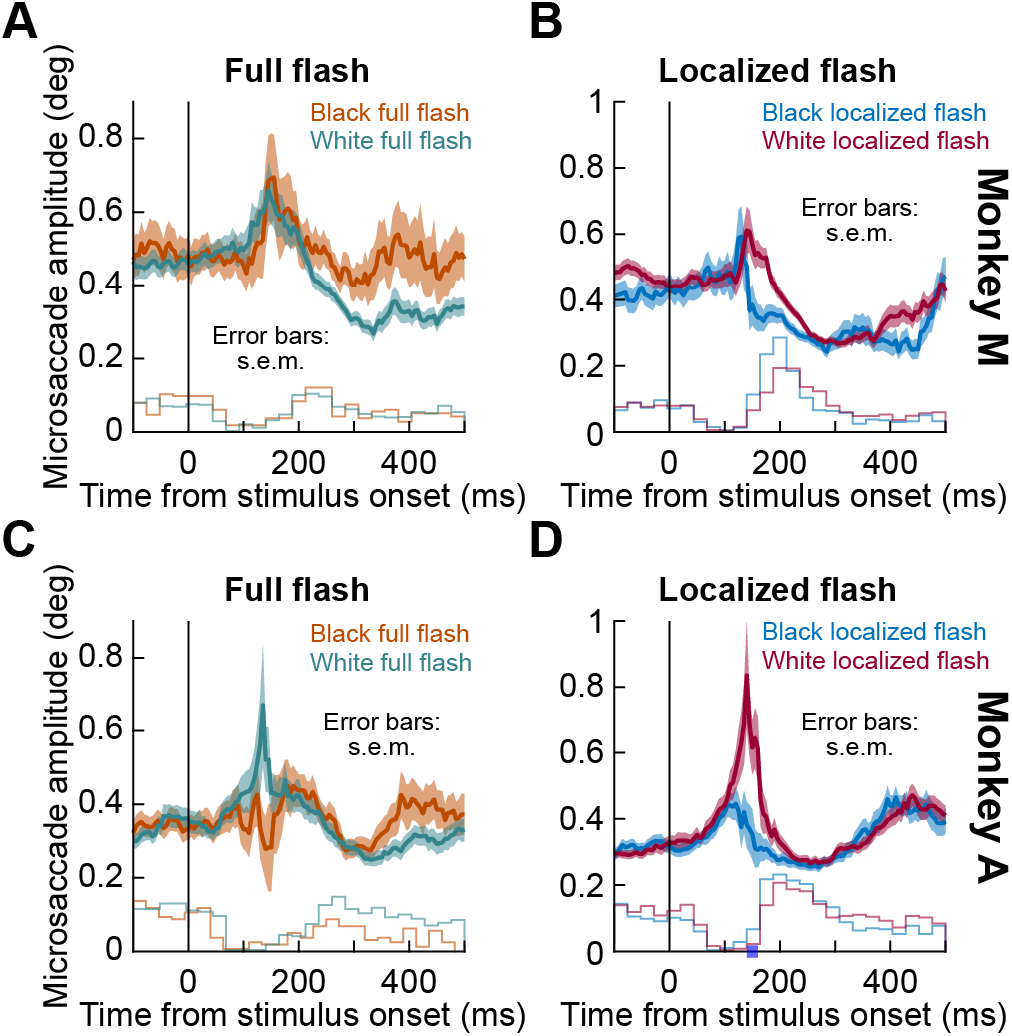
Effects of white and black flashes on microsaccade amplitude time courses with diffuse and localized stimuli. For each monkey (rows) and each flash size (columns), we compared microsaccade amplitude time courses with white or black flashes. There was no strong dependence on stimulus polarity in microsaccade amplitudes. The blue label on the x-axis in **D** indicates the only significant cluster of a short-lasting negative mean difference between black localized and white localized flashes (at 140-160 ms, p = 0.031; cluster-based permutation test). The histogram conventions are the same as in Fig. 6. All other conventions are similar to Fig. 1.

### Microsaccade amplitudes are correlated with directional biases in microsaccades after localized flashes, whether white or black

Finally, to directly test the link between the amplitude modulation caused by our localized stimuli and microsaccade directions, we analyzed the time courses of microsaccade amplitudes with localized flashes, now split based on microsaccade congruency for both black (Fig. 9A, C) and white (Fig. 9B, D) flashes. In both monkeys, the amplitude of congruent saccades was modulated by stimulus onset and increased relative to the pre-stimulus period in the early inhibition phase, as predicted by previous findings (Hafed and Ignashchenkova 2013), thereby confirming our inferences in relation to Fig. 7. Cluster-based permutation tests revealed that this effect was significantly different from the amplitude of incongruent microsaccades in black flashes for monkey M (Fig. 9A) and in white flashes in monkey A (Fig. 9D) (p = 0.009 for monkey M and p = 0.004 for monkey A). Monkey M also demonstrated a similar trend for white flashes (Fig. 9B). Permutation tests run on average amplitudes in the time window of +/-5 ms around the maximum inhibition rates did not reveal additional differences: in fact, only in the black flash condition, the amplitude of congruent saccades was significantly larger than that of incongruent ones (mean difference = 0.406 deg, p = 0.001) in monkey M. Consistent with our analysis of the microsaccade amplitude modulation by the stimulus polarity (Fig. 8), this effect did not depend on whether the flash was black or white.

**Figure 9.**
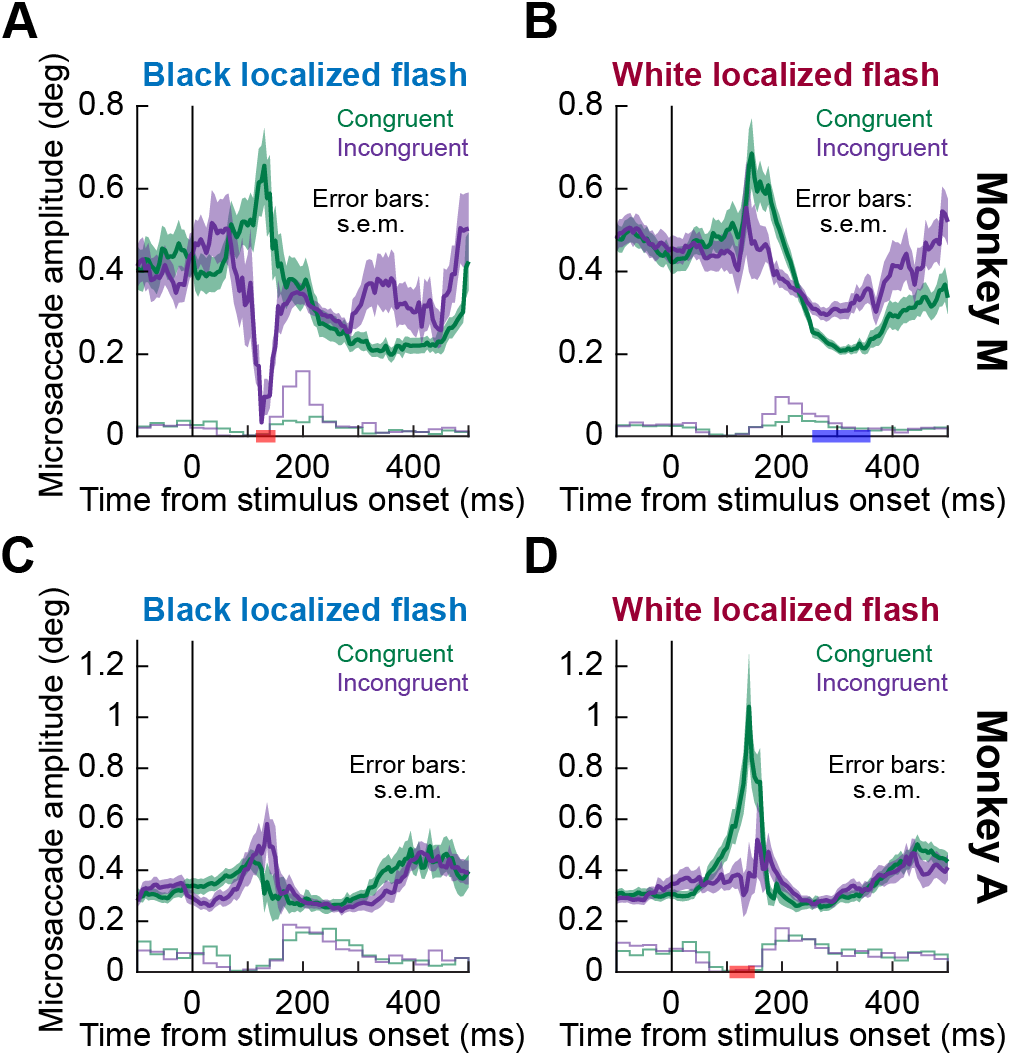
Time courses of microsaccade amplitudes with black and white localized flashes when separated based on microsaccade direction. Both black and white localized flashes caused an increase in the amplitude of microsaccades directed towards the flash (green curves) during the initial inhibition phase of the microsaccadic rate signature, and its difference with the amplitude of incongruent microsaccades became significant in white flashes for monkey A (indicated by the red horizontal label in **D**) and black flashes for monkey M (indicated by the red horizontal label in **A**). Monkey M also demonstrated a similar trend for white flashes (**B**). During the later post-inhibition phase, the amplitude of both congruent and incongruent microsaccades returned to or went slightly below the baseline, and the pattern did not depend on microsaccade direction, with the exception of the white localized flashes in monkey M, where the amplitude modulation of congruent microsaccades was stronger by the end of the rebound period (indicated by the blue label on the x-axis in **B**). The histograms at the bottom of each panel show, in the corresponding color, the frequency of congruent and incongruent microsaccades under a given condition. The histogram conventions are the same as in Fig. 6. All other conventions are similar to Fig. 1.

In contrast, during the later post-inhibition phase, microsaccade amplitudes decreased and went back to, or even below, the pre-stimulus baseline; this was true for both congruent and incongruent microsaccades and did not depend on stimulus polarity, except for the white localized flash condition in monkey M (Fig. 9B), where the amplitude of microsaccades directed towards the stimulus dramatically decreased as compared to incongruent microsaccades at 250-360 ms after stimulus onset (p = 0.016, cluster-based permutation test; i.e. by the end of the rebound phase). No differences were found in the average amplitudes in the time window of +/-5 ms around the peak rebound rates for either monkey, again with the exception of smaller congruent saccades for white flashes in monkey M (mean difference = −0.077 deg, p = 0; permutation test).

Therefore, all of the above results taken together suggest that the strongest overall effects of stimulus polarity emerged with small, localized flashes for which microsaccade rate in the rebound phase of the microsaccadic rate signature was the strongest with black, rather than white, flashes. This was associated with related effects on microsaccade directions and amplitudes. With diffuse flashes, black stimuli were as effective as white ones, in general, whether on microsaccadic inhibition or subsequent rebound.

## Discussion

We investigated the effects of stimulus polarity and size on the microsaccadic rate signature after stimulus onsets. We exploited the fact that even subtle and highly fleeting flashes of only ~8 ms duration are sufficient to cause rapid microsaccadic inhibition after their occurrence followed by a rebound in microsaccade rate. We found that the inhibition was similar for small, localized flashes and large, diffuse ones. However, the subsequent rebound was completely absent with the latter flashes. In terms of stimulus polarity, it had the biggest effects with localized flashes. For these localized flashes, black stimuli caused more substantial changes in the microsaccadic rate signature than white ones, and particularly in the rebound phase after the initial microsaccadic inhibition had ended.

Our results can inform hypotheses about the neural mechanisms for microsaccadic and saccadic inhibition. In (Hafed and Ignashchenkova 2013), we hypothesized that the rate signature reflects visual neural activity in oculomotor areas like, but not exclusively restricted to, the SC. We specifically hypothesized that the dissociation between rate and direction effects (also present in our own data; e.g. Fig. 6) might reflect spatial read out of SC visual activity for the direction effects (Buonocore et al. 2017a) but additional, and potentially different, use of visual activity by the oculomotor system to inhibit saccades for the rate effects (Hafed and Ignashchenkova 2013). Consistent with this, in our current experiments, the similarity that we observed for microsaccadic inhibition between small and large stimuli (Fig. 1) suggests that the early rate effect (i.e. microsaccadic inhibition) is an outcome of early sensory activity that is not necessarily strictly spatial in organization. We hypothesized earlier (Hafed and Ignashchenkova 2013) that a candidate area for realizing such rapid saccadic inhibition could be a late motor area with access to early sensory information. Our ongoing experiments in our laboratory, comparing V1, SC, and brainstem omnipause neurons (Buttner-Ennever et al. 1988; Everling et al. 1998; Gandhi and Keller 1999), strongly support the hypothesis that it is visual sensation by omnipause neurons that is most likely to mediate saccadic inhibition (Buonocore et al. 2020). This would be consistent with our present observations on similar inhibition between small and large stimuli.

The difference in post-inhibition microsaccadic rebound that we observed between small and large stimuli is also consistent with spatially-organized maps for the spatial components of saccadic inhibition (Buonocore et al. 2017a; Hafed and Ignashchenkova 2013). Specifically, with localized flashes, spatial read out of visual stimulus location, say in SC, would cause direction oscillations of microsaccades (Tian et al. 2016). On the other hand, diffuse stimuli centered on fixation would activate symmetric populations of neurons simultaneously. This might not “attract” early microsaccades in any one direction and therefore alleviates the need for opposite microsaccades to occur later in the post-inhibition microsaccadic rebound phase. Thus, with diffuse and symmetric flashes, early microsaccades near the inhibition phase would not introduce large foveal eye position errors like might happen with small, localized peripheral cues. As a result, there would be no need to trigger corrective microsaccades after the inhibition. Indeed, in our earlier work, we showed that shaping the landscape of peripheral visual activity in an oculomotor map, either with extended bars or with simultaneous stimulus onsets at multiple locations, not only influences the directions of early microsaccades, but it also affects subsequent post-inhibition microsaccades, which become oppositely directed from the earlier ones (Hafed and Ignashchenkova 2013). Moreover, we later confirmed that eye position error was indeed an important factor in whether microsaccades were triggered or not (Tian et al. 2018; 2016). Naturally, in behaviors like reading, in which the subsequent forward saccade after any flash is a necessity imposed by the behavioral task at hand, full-screen flashes would be expected to exhibit some post-inhibition rate rebound. This was shown previously (Reingold and Stampe 2004), although even in that study, rebound rates were higher with localized flashes.

Concerning stimulus polarity itself, it is very intriguing that its largest effects appeared on the post-inhibition rebound phase after small, localized flashes (Figs. 3, 5). In our earlier models, we had modeled post-inhibition microsaccades as being driven with greater “urgency” than baseline microsaccades (Hafed and Ignashchenkova 2013; Tian et al. 2016) as if there is extra drive associated with them, needed to recover from the disruptions caused by the stimulus onsets. In later experiments, when we reversibly inactivated FEF, we found that the greatest effects on the microsaccadic rate signature were on post-inhibition microsaccades (Peel et al. 2016), suggesting that the extra drive might come from frontal cortical areas. This might make sense in retrospect: while inhibition may be mediated by rapid, reflexive responses of the oculomotor system to sensory stimulation, post-inhibition eye movements might reflect processes attempting to recover from external disruptions to the ongoing oculomotor rhythm. These processes likely involve additional drive from cortex, a suggestion also made for large saccades (Buonocore et al. 2017b). Our current results of differential effects of black localized stimuli on post-inhibition microsaccades add to the evidence above that different components of the microsaccadic rate signature (e.g. inhibition versus rebound) are governed by distinct and dissociable neural mechanisms.

Concerning why or how stimulus polarity revealed the differences alluded to above, we think that lags between black and white flashes during inhibition (e.g. Figs. 3, 5) might reflect the differences in time that it takes to propagate visual information from the retina to other structures for dark versus light stimuli. For example, it was shown that darks propagate faster than lights to visual cortex due to functional asymmetries in ON and OFF visual pathways (e.g. Westner and Dalal 2019) – in humans; in cats: Jin et al. 2011; Jin et al. 2008; Komban et al. 2014), although it is not absolutely clear at which level of the visual system the temporal advantages of darks first emerge. On the other hand, it is interesting that in our case, black stimuli enhanced microsaccadic rebound rates with localized flashes without necessarily affecting the timing of the rebound microsaccades so much. So, it is not just a matter of speed of the visual pathways that may be at play. An additional factor could be that the visual system might be more sensitive to lower luminances irrespective of the contrast (e.g. in texture discrimination tasks; Chubb and Nam 2000). In addition, we have to consider that our black localized stimuli had more contrast relative to the fixation spot than the white stimuli (although the black flashes were spatially far from the fixation spot, so this effect of contrast relative to the fixation spot might not be so critical). For larger saccades during reading (Reingold and Stampe 2003), black flashes seemed to cause stronger inhibition, but the problem there is that their white flashes did not occlude the black text; thus, their white flashes were likely lower in contrast than their black flashes.

Regardless of the exact causes, our results on stimulus polarity might also be relevant for attention studies since microsaccades are often described as a biomarker for attentional shifts (Engbert and Kliegl 2003; Hafed and Clark 2002; Pastukhov and Braun 2010; Tian et al. 2018; 2016). Our results can therefore allow making predictions with respect to cueing effects demonstrated in typical Posner-style cueing tasks (Posner 1980; Posner and Cohen 1984). For example, our observations might partially explain the mixed results for cue luminance manipulations in cueing paradigms. In these manipulations, varying the cue luminance energy is usually coupled with varying stimulus contrast relative to the background (e.g. Hughes 1984; Mele et al. 2008; Wright and Richard 2003; Zhao and Heinke 2014). Thus, dark cues are necessarily perceptually degraded when compared to bright cues, because of their reduced contrast levels relative to the background. This means that it is still not entirely clear how stimulus polarity factors in cueing paradigms. In our study, the post-inhibition rate of incongruent microsaccades was significantly higher with black localized stimuli than with white localized ones. Taking into account that inhibition of return (IOR) (Klein 2000; Posner and Cohen 1984), which occurs exactly during the post-inhibition phase, might be a direct outcome of the increased likelihood of incongruent microsaccades (Hafed et al. 2015; Tian et al. 2018; 2016), we would predict a stronger IOR effect caused by dark cues as opposed to white cues but no pronounced effect of stimulus polarity on the early facilitation component, when all other experimental parameters such as the relative contrast of white and black stimuli are kept at comparable levels.

## Acknowledgements

We were funded by the Deutsche Forschungsgemeinschaft (DFG) through the Research Unit: FOR 1847 (project: HA6749/2-1). We were also funded by the Werner Reichardt Centre for Integrative Neuroscience (CIN; DFG EXC307) and the Hertie Institute for Clinical Brain Research. TM and ZMH were additionally supported by a CIN intramural grant (Mini_GK 2017-04).

## Author contributions

AB and ZMH collected the data. TM, AB, and ZMH analyzed the data. TM, AB, and ZMH wrote and edited the manuscript.

